# A Sandwich-Structured Silk Fibroin Mesh with ROS-Responsive and Immunoregulatory Functions for Pelvic Floor Repair

**DOI:** 10.64898/2026.05.27.728119

**Authors:** Zhuowei Shen, Yanjiu Li, Xianying Chen, Dehao Tuo, Yaqin Li, Mao Tang, Shiyan Wang, Bing Xiao, Jiaqi Wang, Guikang Wang, Xiaotong Wu, Yue Zhang, Shiwen Zheng, Xuyun Huang, Di jia, Xiuli Sun, Jianliu Wang

## Abstract

Current biodegradable meshes for pelvic floor repair are constrained by a limited ability to actively modulate the hostile immune microenvironment following implantation. To address these challenges, we developed a functionalized degradable silk fibroin mesh (SFM) integrated with a reactive oxygen species (ROS)-responsive nanocomposite hydrogel. The resulting composite mesh, SFM@Gel-NP, features a “hydrogel–mesh–hydrogel” sandwich structure, wherein the hydrogel layers are loaded with self-assembled nanoparticles (NP-H-CRFP) for the co-delivery of the STING inhibitor H–151 and the ROS–scavenging agent catechin. This design provides immediate mechanical reinforcement while enabling microenvironment–triggered drug release. In vitro, NP-H-CRFP demonstrated efficient cellular uptake, significant ROS clearance, and effective attenuation of macrophage inflammation and apoptosis. In vivo, SFM@Gel-NP remodeled the local immune milieu by inhibiting the ROS/cGAS–STING/NF–κB axis, thereby promoting a shift from pro–inflammatory M1 toward pro–regenerative M2 macrophage polarization. This immunomodulatory effect, coupled with enhanced and well–organized collagen deposition—particularly of early type III collagen—resulted in improved tissue integration and repair. This work presents a novel strategy that combines structural reinforcement with active immune regulation, offering a promising next–generation solution for durable and functional pelvic floor reconstruction.

## Introduction

Pelvic floor repair mesh implantation is a primary surgical intervention for severe pelvic organ prolapse (POP)^1, 2^. Currently, clinical meshes are categorized into non-degradable and degradable types. Non-degradable meshes, represented by polypropylene, provide lasting mechanical support but frequently elicit a chronic foreign body reaction (FBR) post-implantation. This often leads to severe complications such as mesh contraction, exposure, erosion, and chronic pain, sometimes necessitating surgical removal^3, 4^. Due to these risks, the U.S. FDA prohibited the transvaginal implantation of synthetic mesh for pelvic floor repair in 2019^5^. Although not entirely banned in China, its clinical application is strictly regulated. Consequently, research focus has shifted towards degradable biomaterials like porcine small intestinal submucosa (SIS) and acellular dermal matrix (ADM)^6–8^. However, these materials are confronted with a dual challenge: not only insufficient long-term mechanical support, but also a lack of inherent active immunomodulatory capacity, which can result in excessive fibrosis or aberrant degradation, ultimately compromising the repair outcome^9^. Therefore, developing novel degradable biomaterials that combine adequate mechanical properties with excellent biocompatibility, and further functionalizing them to actively regulate host immunity (e.g., by loading anti-inflammatory agents or modulating macrophage polarization), has become a key research focus and challenge in pelvic floor reconstruction^10^.

Silk fibroin, a natural degradable polymer, offers unique advantages including abundant sources, tunable mechanical strength, and good biocompatibility, and is already widely used clinically as surgical suture^11^. It can be processed into various forms—fibers, films, hydrogels, and porous scaffolds—catering to diverse biomedical needs and providing a versatile platform for tissue regeneration, drug delivery, and medical device development^12, 13^. Despite its overall favorable biocompatibility, our preliminary studies indicate that silk-based implants can still trigger excessive or prolonged inflammatory responses in the early post-implantation phase^14^, posing a significant obstacle to long-term integration and functional reconstruction. Thus, regulating the local immune response and maintaining immune microenvironment homeostasis following silk mesh implantation is a critical issue that must be addressed to advance its clinical translation.

Reactive oxygen species (ROS), such as superoxide anion (O₂⁻), hydrogen peroxide (H₂O₂), and hydroxyl radicals (·OH), are highly reactive molecules. Biomaterial implantation induces an inflammatory response that elevates local ROS levels, leading to oxidative stress^15–17^. This activates pathways like NF-κB, promoting the release of pro-inflammatory cytokines and potentially resulting in chronic inflammation. Persistent inflammation not only hinders tissue regeneration but may also cause fibrotic encapsulation, compromising implant integration and function^18^. Furthermore, ROS-induced oxidative DNA damage can lead to the release of mitochondrial DNA (mtDNA) into the cytoplasm, activating the cGAS-STING signaling pathway and exacerbating tissue injury^19^. Consequently, an ideal biomaterial should possess anti-ROS capabilities to mitigate oxidative stress and DNA damage, while also inhibiting the cGAS-STING pathway to prevent the amplification of inflammatory cascades^20^.

Herein, we designed and synthesized a copolymer, CRFP, using thioketal and catechin as functional building blocks to confer ROS-responsive properties (Sheme 1A). The thioketal linkages in the polymer backbone undergo specific cleavage in ROS-rich microenvironments, while the pendant catechins provide ROS-scavenging functionality via their polyphenolic hydroxyl groups, synergistically modulating the inflammatory milieu^21, 22^. A nano-delivery system (NP-H-CRFP) was developed via self-assembly to encapsulate the STING inhibitor H-151, and its ROS-triggered release profile was validated using a dialysis method (Sheme 1B). Furthermore, a sandwich-structured composite mesh, termed SFM@Gel-NP, was constructed by laminating NP-H-CRFP-loaded silk fibroin hydrogel onto both sides of a silk fibroin mesh, resulting in a “hydrogel-mesh-hydrogel” architecture that integrates controlled drug release with mechanical compatibility (Sheme 1C). The physicochemical properties of the composite mesh were systematically characterized using rheology and scanning electron microscopy (SEM). *In vitro* cellular assays, including confocal laser scanning microscopy (CLSM) and flow cytometry, demonstrated its anti-inflammatory and anti-apoptotic effects. Subsequently, animal studies were conducted to verify the *in vivo* biocompatibility and safety of the mesh, and to elucidate its mechanism of action—namely, alleviating inflammation and promoting repair by scavenging ROS and inhibiting the cGAS-STING/NF-κB pathway. In summary, we have developed an innovative nanoparticle delivery system and a sandwich-structured mesh that simultaneously targets the ROS and cGAS-STING pathways (Sheme 1D). This strategy significantly enhances the efficacy of anti-inflammatory therapy while ensuring the requisite mechanical performance of the implant *in vivo*, offering a novel and promising solution for the precise treatment of pelvic organ prolapse.

**Scheme 1.**
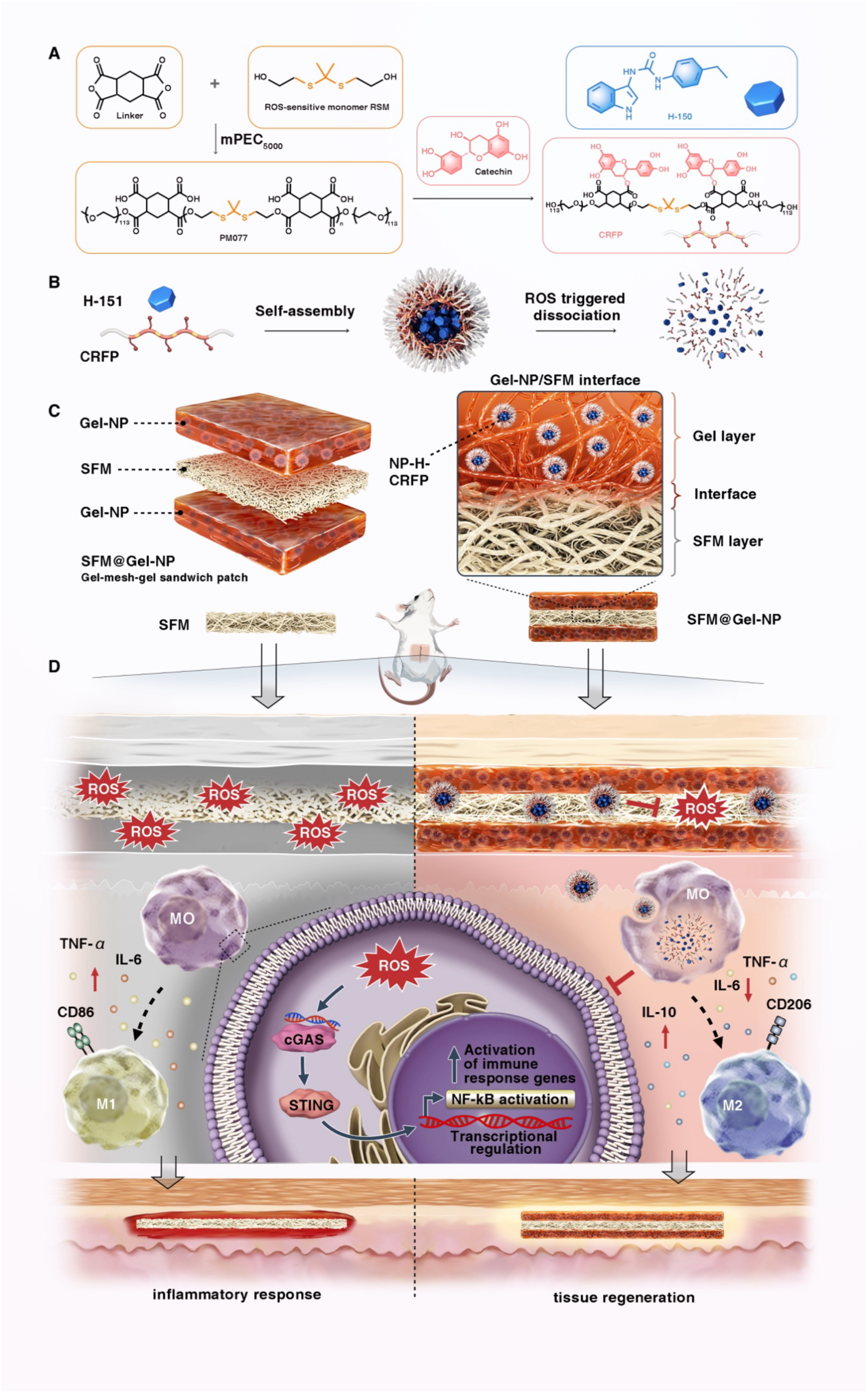
Schematic Illustration of the ROS-Responsive and Immunomodulatory Composite Mesh for Pelvic Floor Repair. (A) Design and synthesis of the ROS-responsive copolymer CRFP, incorporating thioketal (ROS-cleavable) linkages and pendant catechins (ROS-scavenging) for microenvironment modulation. (B) Self-assembly of CRFP with the STING inhibitor H-151 to form nanoparticles (NP-H-CRFP). (C) Fabrication of the sandwich-structured composite mesh (SFM@Gel-NP) by integrating NP-H-CRFP-loaded silk fibroin hydrogel onto both sides of a silk fibroin mesh. (D) Overall therapeutic strategy: the implanted SFM@Gel-NP alleviates inflammation and promotes tissue repair by scavenging ROS and inhibiting the cGAS-STING/NF-κB signaling pathway.

## Results and Discussion

### Synthesis and Characterization of CRFP

As illustrated in Figure 1A, the reactive oxygen species (ROS)-responsive polymer PM077 was synthesized following a previously established protocol (Figure S1)^23^. The target polymer, CRFP, was subsequently obtained by conjugating catechins to the PM077 backbone. The chemical structure of CRFP was confirmed by ¹H nuclear magnetic resonance (¹H-NMR) spectroscopy and Fourier-transform infrared (FTIR) spectroscopy (Figures S1 and S2). Notably, gel permeation chromatography (GPC) analysis revealed a shift in the elution time of CRFP from 13.8 min to 22.2 min after 24-hour incubation with 10 mM H₂O₂, indicating oxidative cleavage of the polymer backbone. This result demonstrates that the incorporated ROS-sensitive monomer (RSM) within the CRFP chain confers specific ROS-responsive properties to the polymer^24^.

**Figure 1.**
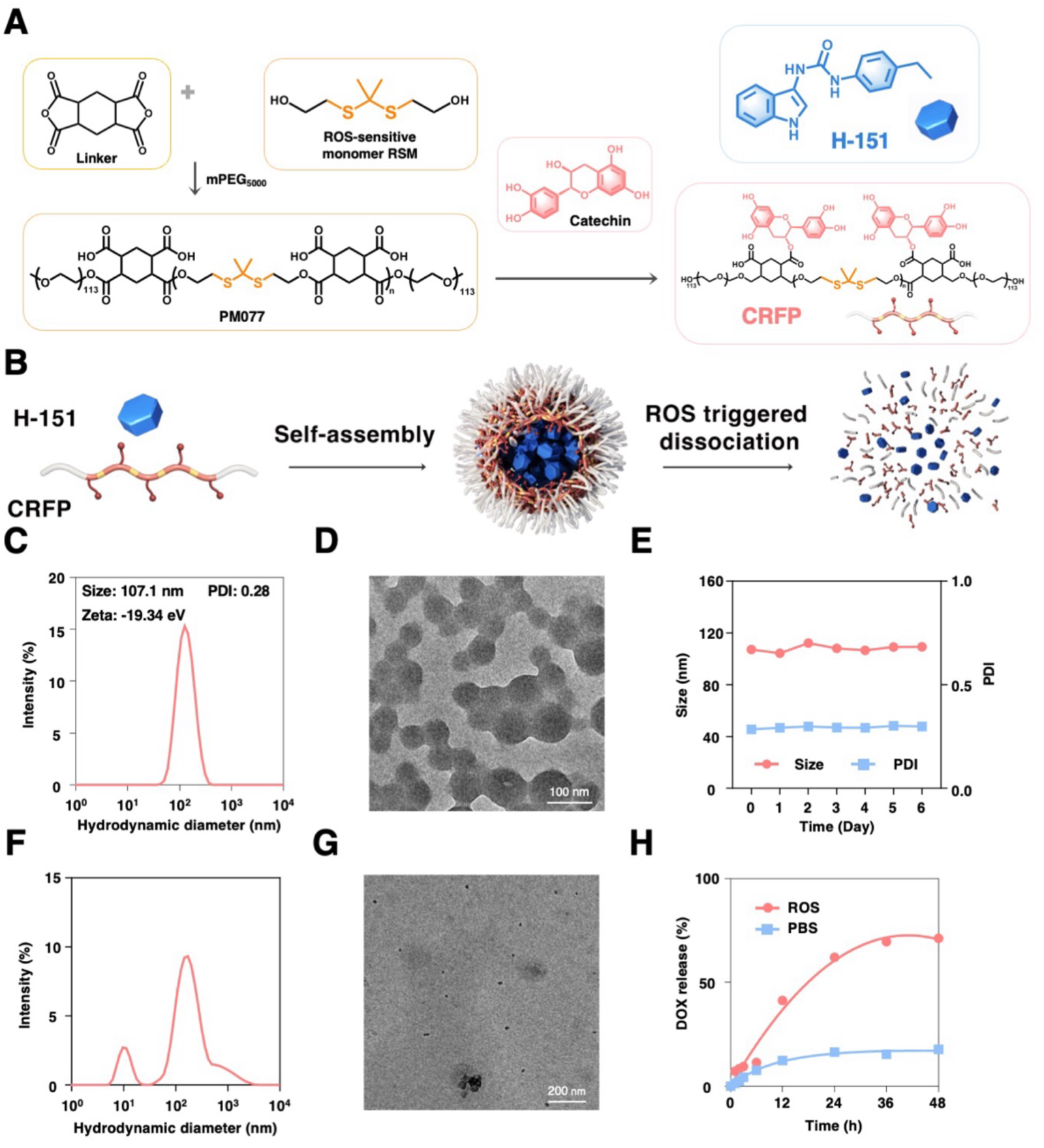
Construction and characterization of NP-H-CRFP. (A) Synthetic route of the CRFP polymer. (B) Schematic illustration of NP-H-CRFP preparation and ROS-responsive disassembly. (C) Hydrodynamic diameter, PDI, and ζ-potential of NP-H-CRFP determined by DLS. (D) TEM image of NP-H-CRFP. Scale bar: 100 nm.(E) Changes in hydrodynamic diameter and PDI of NP-H-CRFP in PBS over 6 days.(F) Hydrodynamic size distribution of NP-H-CRFP after treatment with 10 mM H₂O₂ for 24 h.(G) TEM image of NP-H-CRFP after treatment with 10 mM H₂O₂ for 24 h.(H) In vitro drug release profile of NP-DOX-CRFP under 10 mM H₂O₂ conditions, with DOX used as a model payload.

### Preparation and Characterization of NP-H-CRFP Nanoparticles

Nanoparticles (NP-H-CRFP) were prepared by co-assembling the amphiphilic polymer CRFP with the STING antagonist H-151^25^. Dynamic light scattering (DLS) analysis indicated that the NPs had an average hydrodynamic diameter of 107.1 nm with a narrow size distribution (polydispersity index, PDI = 0.28) and a zeta potential of −19.34 mV (Figure 1C). Their spherical morphology and uniform size (approximately 98 nm in diameter) were further confirmed by transmission electron microscopy (TEM) (Figure 1D). The NPs demonstrated high colloidal stability, maintaining a size variation of less than 10% (within 120 nm) and a PDI below 0.3 over 6 days at 4 °C (Figure 1E).

We next evaluated the ROS-responsive disassembly and drug release behavior of NP-H-CRFP. Upon incubation with 10 mM H₂O₂ for 24 hours, DLS analysis revealed a pronounced change in the particle size distribution. TEM imaging corroborated this finding, showing a clear disruption of the initial spherical nanostructure. Using doxorubicin (DOX)-loaded NPs (NP-DOX-CRFP) as a model system, drug release profiles were assessed. As shown in Figure 1H, over 70% of DOX was released within 48 hours under 10 mM H₂O₂ conditions, a rate significantly accelerated compared to the 18% release observed in PBS. These results collectively confirm the robust ROS-responsive degradation and cargo release capability of the NP-H-CRFP formulation.

### In Vitro Cellular Uptake and Cytoprotective Effects of NP-H-CRFP

To evaluate cellular internalization, NP-H-CRFP was labeled with the near–infrared fluorophore Cy5.5 (forming NP–Cy5.5–CRFP) and incubated with RAW 264.7 macrophages for varying durations. Uptake was quantified using confocal laser scanning microscopy (CLSM) and flow cytometry. CLSM images revealed a time–dependent increase in intracellular fluorescence, indicating efficient and progressive cellular accumulation of the nanoparticles (Figure 2A)^26^. Flow cytometry further confirmed this trend, showing that the mean fluorescence intensity in RAW 264.7 cells after 7 h of incubation was approximately 4.5–fold and 2.0–fold higher than that at 1 h and 4 h, respectively (Figures 2B, C).

**Figure 2.**
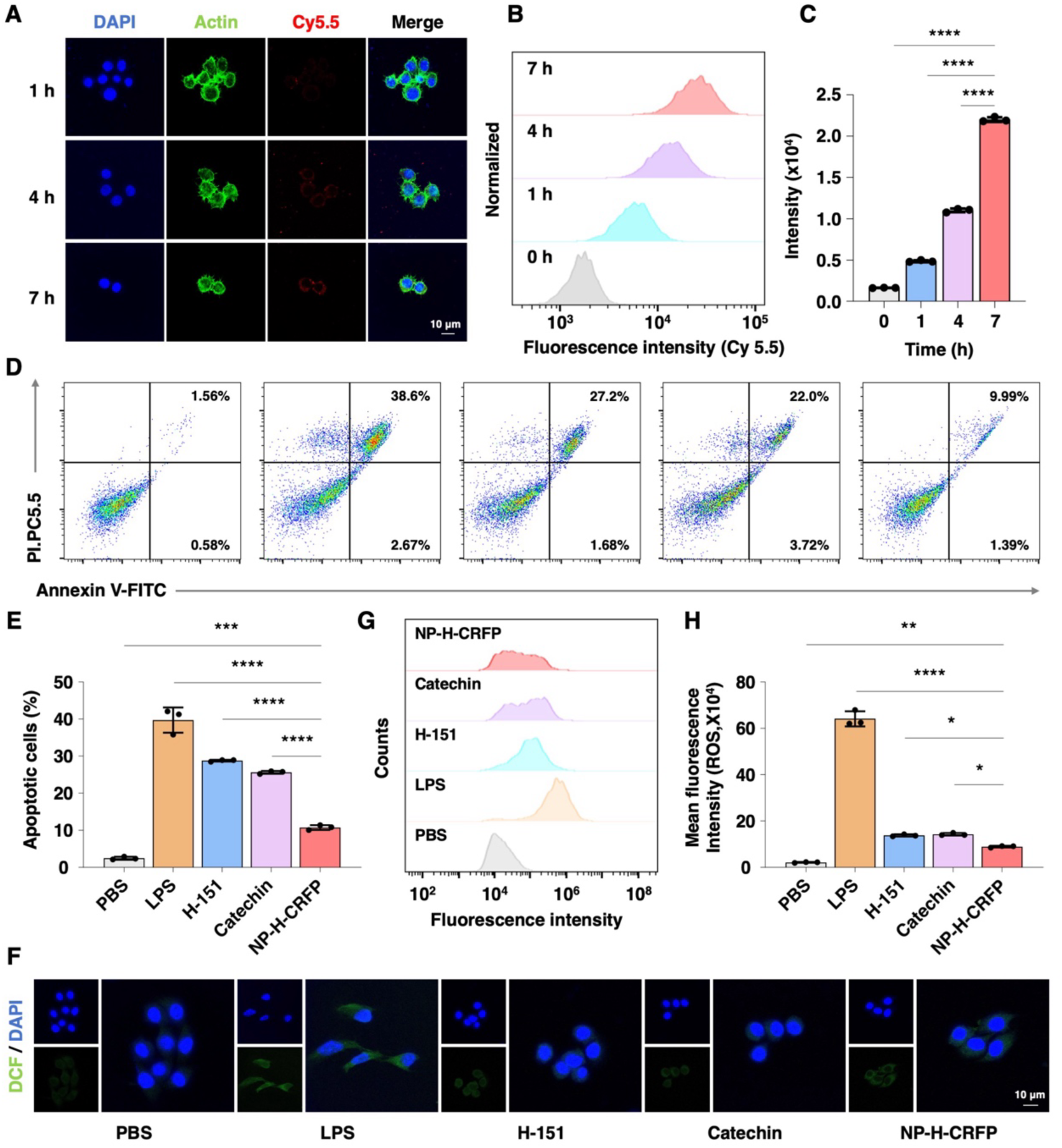
In vitro cellular uptake, anti-apoptotic activity, and ROS-scavenging capability of NP-H-CRFP. (A) Confocal laser scanning microscopy (CLSM) images of RAW 264.7 cells after incubation with NP-Cy5.5-CRFP for 1, 4, and 7 h. Nuclei, cytoskeleton, and NP-Cy5.5-CRFP are shown in blue, green, and red, respectively. Scale bar: 10 μm.(B) Representative flow cytometry profiles showing the internalization of NP-Cy5.5-CRFP by RAW 264.7 cells.(C) Quantitative analysis of NP-Cy5.5-CRFP internalization by RAW 264.7 cells at 0, 1, 4, and 7 h.(D) Representative flow cytometry profiles and (E) quantitative analysis of apoptotic RAW 264.7 cells after different treatments for 24 h.(F) Representative CLSM images showing intracellular ROS levels detected by DCFH-DA staining after different treatments.(G) Flow cytometry analysis of intracellular ROS fluorescence intensity and (H) the corresponding quantitative results. Statistical significance was determined by one-way analysis of variance (ANOVA): p < 0.05, p < 0.01, p < 0.001, p < 0.0001.

Having established cellular uptake, we next assessed the cytoprotective effect of NP–H–CRFP against LPS–induced apoptosis. Flow cytometric analysis showed that stimulation with LPS (20 ng·mL⁻¹) for 24 h markedly increased the apoptosis rate of RAW 264.7 cells. Treatment with free H–151, free catechins, or NP–H–CRFP significantly attenuated this effect. Notably, NP–H–CRFP exhibited the most pronounced anti–apoptotic activity among all treatment groups (Figures 2D, E), indicating that the nano–formulation more effectively alleviates LPS–induced cellular injury.

We further investigated the ability of NP–H–CRFP to modulate intracellular oxidative stress. CLSM and flow cytometry were used to measure ROS levels in RAW 264.7 cells after various treatments. CLSM imaging demonstrated that LPS stimulation strongly elevated intracellular ROS, whereas treatment with H–151, catechins, or NP–H–CRFP reduced ROS signals. The ROS–scavenging effect was most evident in the NP–H–CRFP group (Figure 2F). Quantitative flow cytometry data indicated that the mean fluorescence intensity of ROS in cells treated with free H–151 or free catechins was about 1.5–fold and 1.6–fold higher, respectively, than in cells treated with NP–H–CRFP (Figures 2G, H).

Together, these results demonstrate that NP–H–CRFP is efficiently internalized by macrophages, significantly reduces LPS–induced intracellular ROS accumulation, mitigates oxidative stress, and thereby effectively inhibits apoptosis, highlighting its potent cytoprotective function.

### Preparation and Characterization of the SFM@Gel-NP Composite Mesh

A silk fibroin-based composite hydrogel mesh (SFM@Gel-NP) was constructed as illustrated in Figure 3A. First, a silk fibroin hydrogel was prepared from silk fibroin lyophilized powder, with NP-DOX-CRFP incorporated as a model payload, following a previously reported procedure (Figure 3B). Subsequently, SFM was coated on both sides with this hydrogel to fabricate a “gel–mesh–gel” sandwich-structured mesh, designated SFM@Gel-NP. For comparative evaluation, several control meshes were prepared using the same process: SFM coated with blank hydrogel (SFM@Gel), and sandwich meshes loaded with H–151 or CRFP alone, referred to as SFM@Gel–H151 and SFM@Gel–CRFP, respectively.

**Figure 3.**
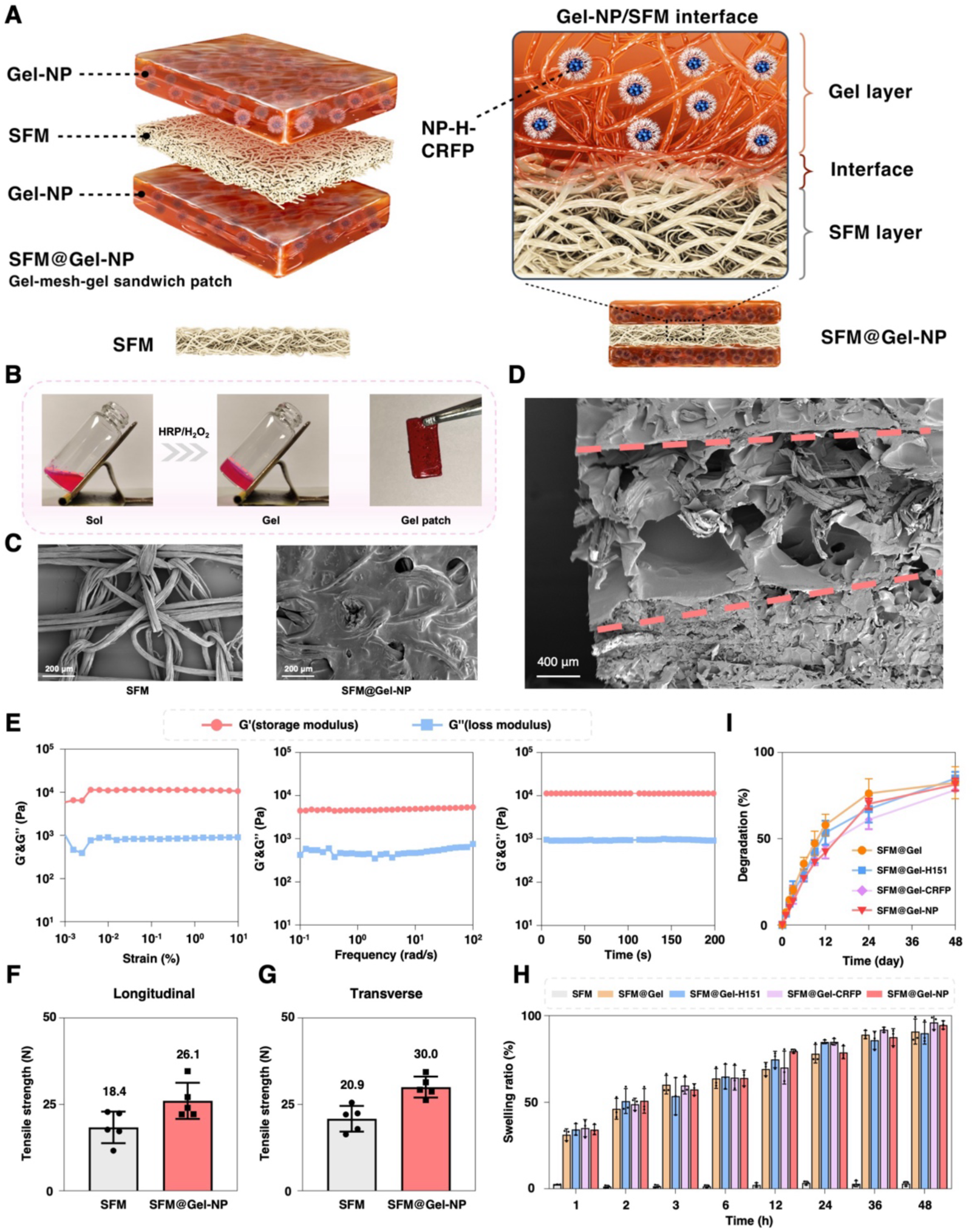
Fabrication and characterization of SFM@Gel-NP. (A) Schematic illustration of SFM@Gel-NP and magnified cross-sectional view of the Gel-NP/SFM interface. (B) Photographs showing HRP/H₂O₂-mediated sol–gel transition and Gel mesh formation. (C) Scanning electron microscopy(SEM) images of SFM and SFM@Gel-NP. Scale bars: 200 μm. (D) Cross-sectional SEM image of SFM@Gel-NP. Scale bar: 400 μm. (E) Rheological characterization of SFM@Gel-NP by strain, frequency, and time sweep tests. (F,G) Longitudinal and transverse maximum tensile loads of SFM and SFM@Gel-NP. (H) Swelling ratios of different meshes. (I) In vitro degradation profiles of different meshes.

Scanning electron microscopy (SEM) comparison of SFM and SFM@Gel-NP confirmed successful hydrogel loading on the mesh (Figure 3C). Cross–sectional SEM imaging further revealed that the silk fibroin hydrogel layers continuously coated both surfaces of SFM and penetrated into the mesh at the interface, forming an interwoven and interpenetrating structure indicative of strong interfacial bonding (Figure 3D and Figure S5)^27^.

Rheological analysis of SFM@Gel-NP via strain, frequency, and time sweeps demonstrated that the storage modulus (G′) consistently exceeded the loss modulus (G″), confirming the formation of a stable, elastic–dominant hydrogel network within the composite system (Figure 3E). Mechanical testing showed that coating with hydrogel significantly enhanced the tensile strength of the mesh. Compared to bare SFM, SFM@Gel-NP exhibited increased maximum tensile forces, from 18.4 N to 26.1 N in the longitudinal direction and from 20.9 N to 30.0 N in the transverse direction (Figures 3F, G), underscoring the reinforcing effect of the hydrogel coating and its strong interfacial integration with the mesh.

The swelling behavior of the composite meshes was assessed by measuring the swelling ratio over time. All hydrogel–containing groups—SFM@Gel, SFM@Gel–H151, SFM@Gel–CRFP, and SFM@Gel–NP—exhibited a gradual increase in swelling ratio, indicating favorable water–absorption capacity (Figure 3H) ^28^. In vitro degradation studies further revealed that these groups could degrade over 70% within 48 days (Figure 3I and Figure S5). Collectively, these results demonstrate that SFM@Gel-NP possesses swelling and degradation profiles comparable to those of the other hydrogel–based meshes, confirming that the incorporation of NP–H–CRFP does not compromise the structural integrity of the fibroin hydrogel network or adversely affect the fundamental stability and degradability of the mesh ^14^.

### Evaluation of the Biosafety and Anti-Inflammatory Activity of the SFM@Gel-NP System

To evaluate the in vivo biocompatibility and anti-inflammatory efficacy of the nanoparticle-loaded, gel-functionalized mesh, five experimental groups were established as illustrated in Figure S8: blank SFM (control), SFM coated with blank hydrogel (SFM@Gel), SFM coated with H–151-loaded hydrogel (SFM@Gel–H151), SFM coated with nanoparticle-loaded hydrogel (SFM@Gel–CRFP), and SFM coated with dual-drug-loaded hydrogel (SFM@Gel–NP). Given the distinct pathogenesis of childbirth-related pelvic floor dysfunction, no universal animal model for pelvic organ prolapse (POP) is currently available. Since pelvic floor repair materials must exhibit both biocompatibility and suitable mechanical properties, the rat abdominal wall defect model has been widely adopted in prior studies ^29–32^.

In this work, twenty-five 8-week-old SD rats were randomly assigned to the five groups (n = 5 per group). Bilateral partial-thickness abdominal wall defects were created as illustrated in Figure 4A, followed by implantation of the corresponding meshes. All animals survived the procedure (25/25) without postoperative complications such as restricted mobility, mesh exposure, or surgical site infection. Tissues were harvested at predetermined time points for analysis. No significant differences in body weight were observed among groups, suggesting that the implants did not induce systemic metabolic disturbances.

**Figure 4.**
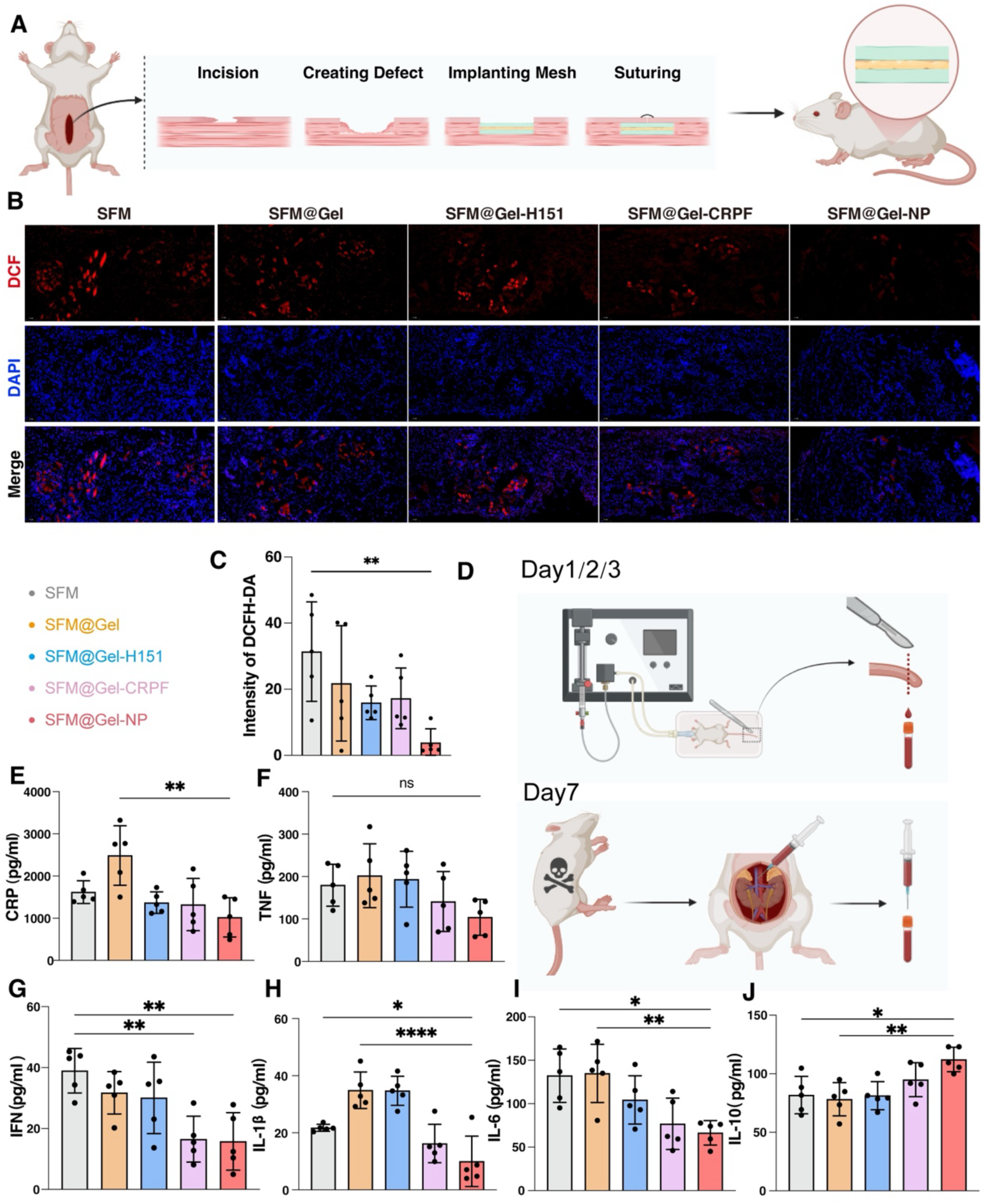
In vivo evaluation of biosafety, local oxidative stress, and systemic inflammatory response following mesh implantation. (A) Schematic illustration of the rat abdominal wall defect model and mesh implantation procedure. (B) Representative fluorescent images of DCFH-DA staining (Red) detecting reactive oxygen species (ROS) in tissues surrounding the implant at day 7 post-surgery. Scale bar, 50 µm. (C) Quantitative analysis of fluorescence intensity from (B). Data are presented as mean ± SD (n = 5). (D) Timeline for blood collection and serum cytokine analysis. (E-J) Serum levels of pro-inflammatory cytokines (IL-1β, CRP, IL-6, IFN-γ, TNF-α) and the anti-inflammatory cytokine IL-10 at postoperative day 2. Data are presented as mean ± SD (n = 5). *P < 0.05, **P < 0.01, ***P < 0.001, ****P < 0.001.

Histological evaluation of major organs (heart, liver, spleen, lungs, and kidneys) via hematoxylin and eosin (H&E) staining is a standard approach to determine whether material degradation products elicit systemic toxicity, thereby confirming overall biosafety^33, 34^. One week after surgery, tissues from the SFM group and the SFM@Gel-NP group were collected for H&E staining and pathological assessment. Compared with controls, the SFM@Gel-NP group showed no notable pathological alterations in major organs, indicating favorable biocompatibility and systemic safety of the functionalized mesh (Figure S9).

Implantation of a mesh elicits a foreign body reaction, a host defense mechanism against non-self materials^35^. This process activates immune cells—including macrophages and neutrophils—which generate reactive oxygen species (ROS). Excessive ROS production promotes oxidative stress and the release of inflammatory cytokines such as TNF-α and IL-6, thereby amplifying local inflammation^36, 37^.

Sustained inflammation can compromise peri-implant tissue integration, cause tissue damage, and contribute to patient discomfort. Our previous in vitro experiments confirmed that nanoparticle uptake suppresses cellular oxidative stress (Figure 2). To evaluate whether the mesh mitigates local oxidative stress in vivo, we performed DCFH-DA staining on tissues surrounding the implant 7 days after surgery. As shown in Figures 4B and 4C, fluorescence intensity was significantly lower in the SFM@Gel-NP group than in the SFM group (P < 0.01), demonstrating that the system effectively attenuates early oxidative stress at the implantation site.

An excessive inflammatory response can disrupt immune homeostasis, potentially leading to a cascade of local or systemic tissue damage accompanied by elevated levels of pro-inflammatory cytokines such as IL-1β, CRP, IL-6, IFN-γ, and TNF-α ^38^. To evaluate whether the implant influences systemic inflammation, we measured serum cytokine levels post-operatively. Blood samples were collected from the tail vein on days 1, 2, and 3, and from the inferior vena cava on day 7 (Figure 4D).

Compared with the SFM and SFM@Gel groups, the SFM@Gel-NP group showed significant downregulation of key pro-inflammatory cytokines across multiple time points: CRP and IL-6 on day 1 (Figure S10); IL-1β, CRP, IL-6, and IFN-γ on day 2 (Figures 4E, 4G–I); IL-1β, CRP, and IL-6 on day 3; and IL-1β, CRP, and TNF-α on day 7 (Figure S10). In parallel, levels of the anti-inflammatory cytokine IL-10, which promotes wound healing^39^, were markedly elevated at all time points in the SFM@Gel-NP group (Figure 4J, Figure S10).

Together, these findings demonstrate that the SFM@Gel-NP composite system not only exhibits favorable biosafety but also effectively mitigates early post-implantation inflammation by suppressing oxidative stress and modulating the expression of key inflammatory cytokines.

### Transcriptomic and Mechanistic Elucidation of Anti-inflammatory and Pro-repair Actions

We first assessed the mechanical integrity of the implanted meshes, as pelvic floor repair materials must withstand tensile stresses exceeding 16 N/cm to tolerate physiological intra-abdominal pressures ranging from 0.2 to 20 kPa, with peaks reaching 13 kPa during coughing^40, 41^. Tensile testing of explanted mesh-tissue complexes one week post-surgery confirmed that all groups met this threshold. Hydrogel loading improved the mechanical properties, with the SFM@Gel-NP group showing an increased maximum tensile strength compared to the SFM group, although the difference was not statistically significant (P > 0.05) (Figures 5A, B).

**Figure 5.**
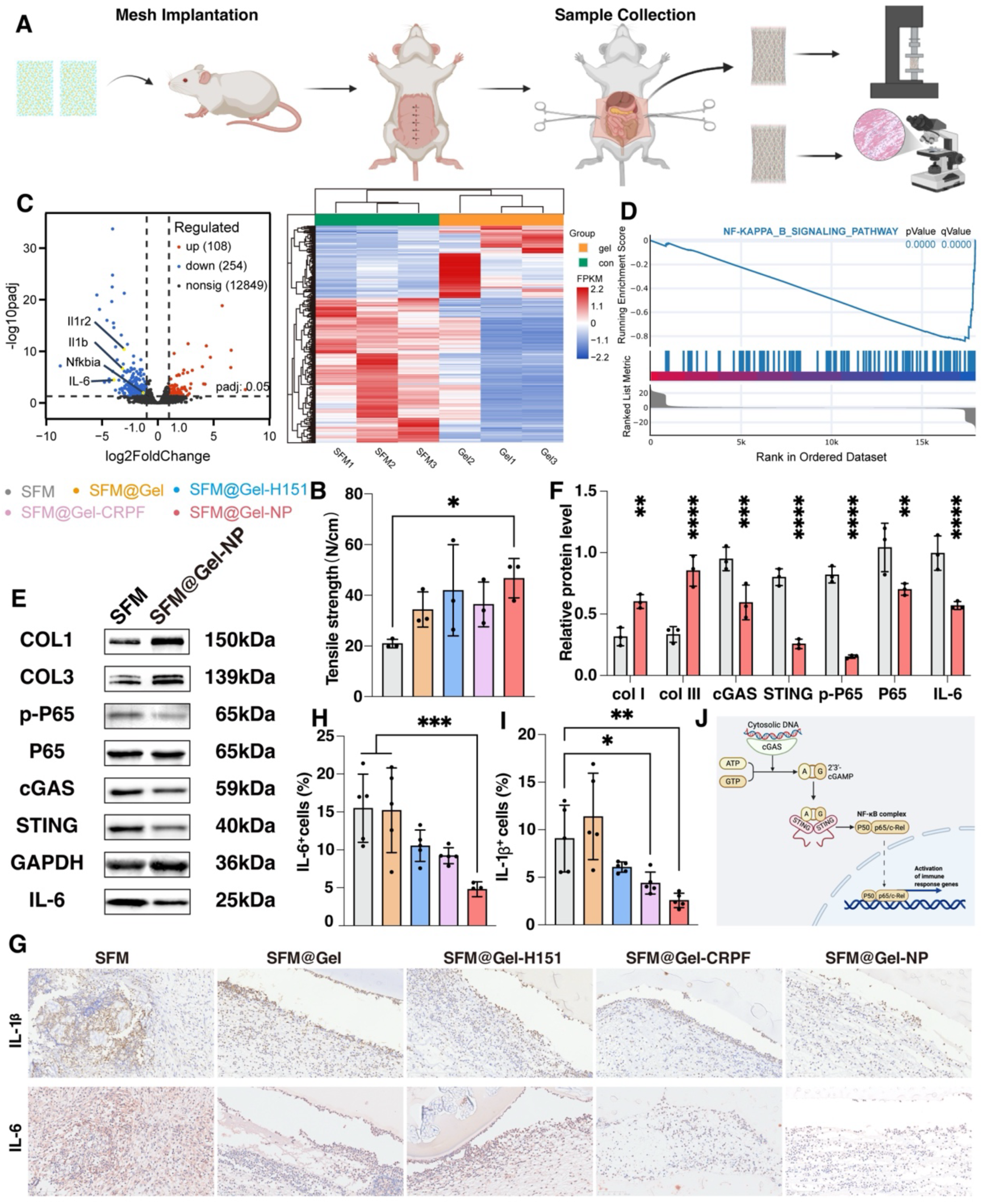
Transcriptomic reveals the anti-inflammatory and pro-repair actions of the functionalized mesh. (A) Schematic illustration of the explanted mesh-abdominal wall tissue complex used for mechanical and molecular analyses. (B) Maximum tensile strength of explanted mesh-tissue complexes from each group at 1-week post-implantation. Data are presented as mean ± SD (n = 3). (C) Volcano plot displaying differentially expressed genes (DEGs) between the SFM and SFM@Gel-NP groups. Red and blue dots represent upregulated and downregulated genes (|log2FC| > 1, p-adj < 0.05), respectively. Key downregulated NF-κB-related genes are labeled. (D) Gene Set Enrichment Analysis (GSEA) plot showing significant suppression of the NF-κB signaling pathway in the SFM@Gel-NP group. (E) Representative western blot images of key proteins from peri-implant tissues, including cGAS, STING, phosphorylated p65 (p-p65), total p65, COL1, COL3, and GAPDH. (F) Densitometric quantification of protein expression levels from (E), normalized to GAPDH. Data are presented as mean ± SD (n = 3). (G-I) Representative immunohistochemical staining images (G) and quantitative analysis (H, I) of IL-1β and IL-6 expression in peri-implant tissues at 1 week. Scale bar, 100 µm. Data are presented as mean ± SD (n = 5). (J) Proposed mechanism: The SFM@Gel-NP system alleviates the post-implantation inflammatory response and promotes repair by inhibiting the ROS/cGAS-STING/NF-κB signaling axis and enhancing collagen deposition. *P < 0.05, **P < 0.01, ***P < 0.005, ****P < 0.001.

To elucidate the molecular mechanism underlying the anti-inflammatory effect, transcriptomic analysis was performed. RNA sequencing of explanted complexes from the SFM and SFM@Gel-NP groups identified 362 differentially expressed genes (DEGs), with 254 downregulated in the SFM@Gel-NP group (Figure 5C). Gene Set Enrichment Analysis (GSEA) revealed the NF-κB signaling pathway as the most significantly suppressed (p = 0.000) (Figure 5D). Consistent with this, key NF-κB-related genes (e.g., *IL-1β*, *IL-6*, *Cxcl1*, *Cxcl2*, *Cxcl3*, *Nfkb*) were downregulated.

Enrichment analyses indicated that the downregulated DEGs were primarily involved in innate immune and inflammatory responses (Figure S11A) and were associated with upstream regulators of the NF-κB pathway, including the PI3K-Akt and TNF signaling pathways (Figure S11B).

Protein-level validation confirmed the inhibition of this regulatory network. Immunofluorescence and western blot analyses demonstrated effective in vivo suppression of the upstream cGAS-STING axis in the SFM@Gel-NP group, as shown by reduced cGAS and STING signals (Figure S12, Figures 5E-F). Furthermore, western blotting confirmed the downregulation of the NF-κB pathway, evidenced by significantly reduced expression of both phosphorylated p65 (p-p65) and total p65—key markers of pathway activation^42, 43^ —in the SFM@Gel-NP group (Figures 5E-F).

Finally, immunohistochemical analysis of peri-implant tissues showed a significant reduction in the local expression of the pro-inflammatory cytokines IL-1β and IL-6 in the SFM@Gel-NP group compared to controls (Figures 5G-I). This protein-level evidence aligns with the transcriptomic data, collectively demonstrating that the SFM@Gel-NP composite system attenuates the inflammatory response through coordinated regulation of the cGAS-STING/NF-κB pathway.

Activation of the cGAS-STING pathway is a known trigger for NF-κB signaling, leading to the production of pro-inflammatory cytokines including IL-6 and TNF-α^44^. Furthermore, reactive oxygen species (ROS) can exacerbate inflammation by activating IκB kinases or inhibiting specific phosphatases, thereby upregulating redox-sensitive NF-κB activity^45^. Integrating our experimental data, we conclude that the SFM@Gel-NP system exerts its anti-inflammatory effect by coordinately inhibiting the ROS/cGAS-STING/NF-κB axis, which in turn suppresses the expression of downstream inflammatory mediators (Figure 5J).

Notably, western blot analysis also revealed an upregulation of collagen synthesis-related proteins, COL1 and COL3, in the SFM@Gel-NP group (Figures 5E-F). This finding suggests that the functionalized mesh not only dampens the inflammatory cascade but may concurrently promote extracellular matrix synthesis and tissue repair, potentially contributing to enhanced mechanical integration and long-term functional recovery at the implantation site.

### Immunomodulatory Effects on Macrophage Polarization and Collagen Remodeling

Macrophages play a pivotal role in the host response to biomaterial implantation, exhibiting plasticity to polarize into pro-inflammatory (M1) or anti-inflammatory/pro-regenerative (M2) phenotypes in response to microenvironmental cues^46^. Following mesh implantation, the generation of reactive oxygen species (ROS) can induce oxidative DNA or mitochondrial damage, releasing double-stranded DNA (dsDNA) that activates the cGAS-STING pathway. This, in turn, triggers the downstream NF-κB pathway. These signaling cascades engage in a positive feedback loop, driving the secretion of pro-inflammatory cytokines like TNF-α and collectively promoting a sustained M1 phenotype^47, 48^. Conversely, scavenging excess ROS and inhibiting the cGAS-STING/NF-κB axis can reduce pro-inflammatory cytokine production and effectively steer macrophage polarization towards the M2 phenotype, which facilitates inflammation resolution and tissue repair through the secretion of factors such as IL-10^49, 50^.

To assess the impact of our meshes on the local immune microenvironment, we performed immunofluorescence staining to identify and quantify macrophage phenotypes using CD68 (pan-macrophage marker), CD86 (M1 marker), and CD206 (M2 marker) (Figure 6A). One week post-implantation, the SFM group showed significant infiltration of CD86⁺ M1 macrophages (yellow), with notably higher numbers than all other groups (Figure 6B). In contrast, the SFM@Gel-CRFP and SFM@Gel-NP groups exhibited a marked increase in the proportion of CD206⁺ M2 macrophages (red), indicating a shift in local macrophage polarization towards an anti-inflammatory and pro-regenerative state (Figure 6C). This transition from a dominant M1 to an enhanced M2 phenotype suggests that the functionalized meshes exert immunomodulatory effects during the critical phase from acute inflammation to tissue repair, thereby reshaping the local microenvironment to favor long-term healing^51^.

**Figure 6.**
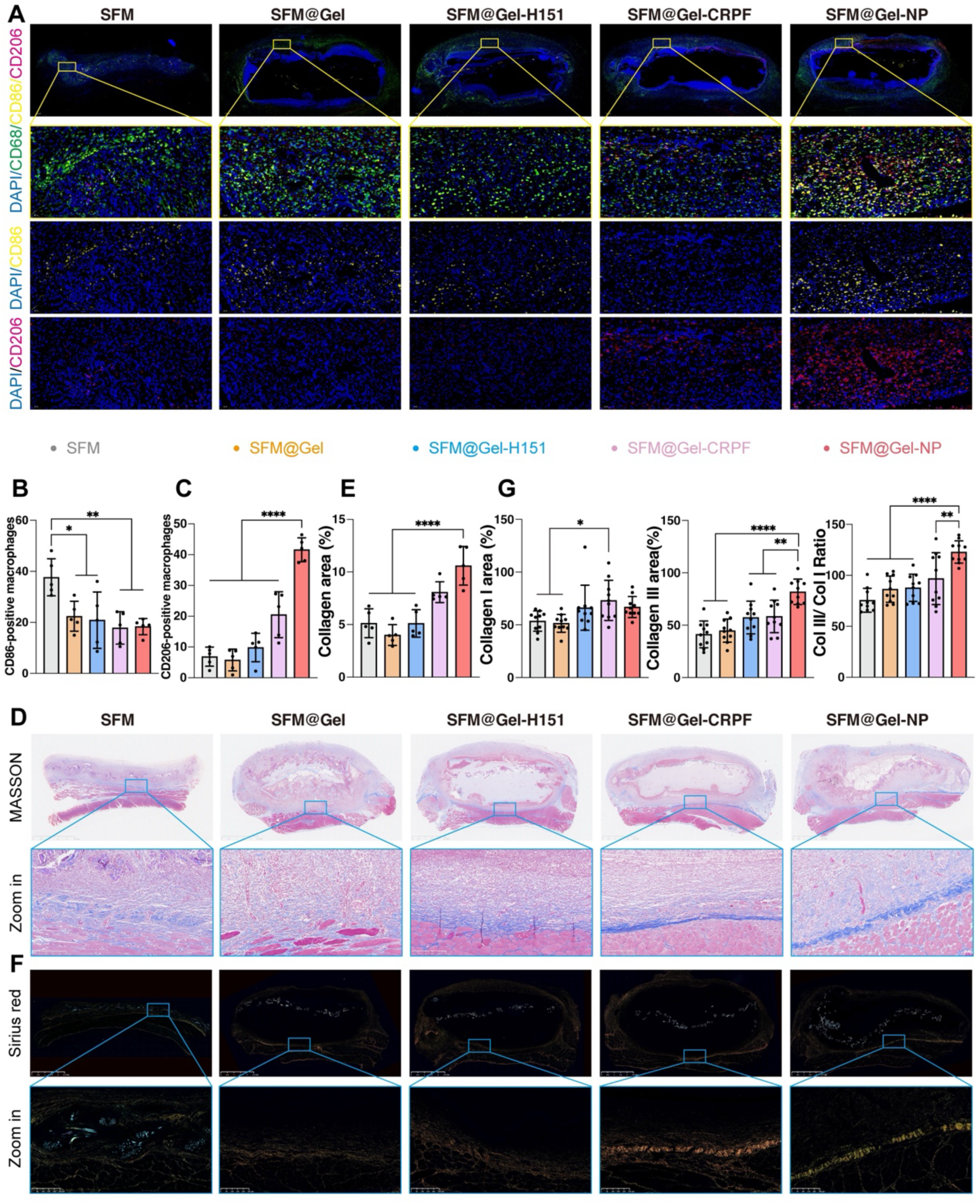
Functionalized mesh modulates macrophage polarization and promotes organized collagen deposition. (A) Schematic diagram of immunofluorescence staining strategy for identifying macrophage phenotypes. CD68 (white) was used as a pan-macrophage marker. Co-localization of CD68 with CD86 (yellow) indicates pro-inflammatory M1 macrophages, while co-localization with CD206 (red) indicates anti-inflammatory/pro-regenerative M2 macrophages.Scale bar, 50 µm. (B) Quantitative analysis of M1 macrophage (CD68⁺CD86⁺) infiltration in peri-implant tissues at 1-week post-surgery.Data are presented as mean ± SD (n = 5). (C) Quantitative analysis of M2 macrophage (CD68⁺CD206⁺) proportion in peri-implant tissues at 1 week. Data are presented as mean ± SD (n = 5). (D) Representative images of Masson’s trichrome staining showing collagen deposition (blue) in the implantation area at 1 week(D). Scale bar, 250 µm. (E) The histogram shows quantitative analysis of collagen density. Data are presented as mean ± SD (n = 5). (F)Representative polarized light microscopy images of Sirius Red-stained sections distinguishing collagen subtypes: mature type I collagen fibers (orange-red) and the immature type III collagen network (green). (G) Quantitative analysis of the area of type I (red birefringence) , type III collagen (green birefringence) and III/I ratio from Sirius Red staining under polarized light. Data are presented as mean ± SD (n = 5). Scale bar, 250 µm. *P < 0.05, **P < 0.01, ***P < 0.005, ****P < 0.001.

To verify the functional consequence of this immunomodulation on tissue regeneration, we evaluated collagen deposition using Masson’s trichrome staining. At one week, the SFM, SFM@Gel, and SFM@Gel-H151 groups displayed only sparse and disorganized collagen fibers (stained blue). In contrast, the SFM@Gel-CRFP and SFM@Gel-NP groups showed dense and well-organized collagen layers, with the most significant collagen regeneration observed in the SFM@Gel-NP group (Figure 6D). Quantitative analysis confirmed that collagen density in the SFM@Gel-NP group was significantly higher than in the first three groups (p < 0.001).

The quality of newly formed collagen is crucial for repair outcomes. Type III collagen, which provides initial, more elastic scaffolding during early wound healing, is later gradually replaced by the stronger type I collagen^52, 53^. Using Sirius Red staining under polarized light, we distinguished collagen subtypes: type I fibers appear orange-red, while the type III network exhibits green birefringence (Figure 6G). Quantitative analysis revealed that the area of green-stained type III collagen was significantly larger in the SFM@Gel-NP group compared to all other groups (Figure 6E, p < 0.01), indicating a repair pattern characterized by pronounced early deposition of type III collagen.

Collectively, these results demonstrate that the SFM@Gel-NP system promotes a favorable tissue repair microenvironment by modulating macrophage polarization from a pro-inflammatory M1 towards a pro-regenerative M2 phenotype. This immunomodulatory effect is functionally linked to enhanced and better-organized collagen deposition, particularly the early accumulation of type III collagen, which supports subsequent tissue remodeling and integration.

## Conclusion

In this study, we developed a functionalized silk fibroin mesh (SFM) for pelvic floor reconstruction, which integrates robust mechanical support with precise immunomodulatory capability. The mesh was engineered with a reactive oxygen species (ROS)-responsive nanocomposite hydrogel, loaded with self-assembled nanoparticles (NP-H-CRFP) for the co-delivery of the STING inhibitor H-151 and the ROS-scavenging agent catechin. This design constitutes a microenvironment-responsive drug delivery system. The resulting composite mesh, SFM@Gel-NP, features a “hydrogel–mesh–hydrogel” sandwich structure that provides immediate mechanical reinforcement while enabling the controlled release of therapeutic agents. SFM@Gel-NP positively remodeled the post-implantation immune microenvironment by inhibiting the ROS/cGAS–STING/NF-κB signaling axis. This intervention promoted a phenotypic shift in macrophages from a pro-inflammatory (M1) state toward a pro-regenerative (M2) state, thereby attenuating chronic inflammation and facilitating superior tissue integration and functional repair. In summary, this work presents a novel biomaterial strategy that successfully combines structural reinforcement with active immunoregulation, addressing both the mechanical and immunological challenges inherent in pelvic floor repair.

## METHODS AND EXPERIMENTS

### Antibodies and reagents

Antibodies against cGas(530423) were purchased from Abmart. Antibodies against STING (19851-1-AP), NF-κB p65(80979-1-RR), IL-1β(26048-1-AP), CD68(83014-5-RR), CD86(26903-1-AP), CD206(18704-1-AP), collagen I (66761-1-Ig) and collagen III (22734-1-AP) were purchased from proteintect. Antibodies against IL-6(ab9324) and Anti-NF-kB p65(ab76302) were purchased from Abcam.

Annexin V-FITCApoptosis Detection Kit(C1062S) was purchased from Beyotime. Reactive Oxygen Species Assay Kit (G1706-100T) was purchased from Servicebio. Cell Counting Kit-8(CK04-500T) was purchased from Dojindo. IL-1β (RX2D302066), IL-6 (RX2D302196), IL-10 (RX302880R), IL-18 (RX302871R), CRP (RX302991R), TNF-α (RX2D310636) and IFN-α (RX302925R) Elisa kits were purchased from Ruixin Biotechnology.

### Synthesis of PM077

PM077 was synthesized by reacting 2,2’-(propane-2,2-diylbis(sulfanediyl))bis(ethan-1-ol) (750 mg) with (S)-ethyl 2,6-diisocyanatohexanoate (1200 mg) in DMF (10 mL) at room temperature for 24 h. Then, mPEG5000 (1200 mg) was added, and the reaction was continued at 50 °C for 24 h. The product was purified by dialysis against deionized water using a dialysis membrane with a molecular weight cutoff of 8000–14000 Da for 2 days, followed by vacuum drying to obtain PM077.

### Synthesis of CRFP

CRFP was synthesized by reacting PM077 with catechin in the presence of EDCI and DMAP at 50 °C for 24 h. The crude product was purified by dialysis against deionized water and then lyophilized to obtain CRFP.

### Preparation of NP-H-CRFP

CRFP (100 mg) and H-151 (10 mg) were dissolved in DMSO (1 mL) and slowly added dropwise into deionized water (10 mL) under stirring for 10 min. The mixture was dialyzed against deionized water using a dialysis membrane with a molecular weight cutoff of 3500 Da to remove DMSO and free H-151. The obtained nanoparticles were denoted as NP-H-CRFP. The H-151 loading content was quantified by high-performance liquid chromatography (HPLC).

### Preparation of NP-DOX-CRFP and NP-Cy5.5-CRFP

NP-DOX-CRFP and NP-Cy5.5-CRFP were prepared using a similar nanoprecipitation method, except that H-151 was replaced with Doxorubicin (DOX) or Cy5.5, respectively. The crude nanoparticle solutions were dialyzed against deionized water to remove organic solvent and unloaded molecules, yielding NP-DOX-CRFP and NP-Cy5.5-CRFP.

### ROS-Accelerated Drug Release from NP-DOX-CRFP

NP-DOX-CRFP solution (10 mL) was transferred into a pre-swelled dialysis bag (MWCO: 3500 Da) and immersed in PBS or H₂O₂-containing PBS (50 mL) under gentle shaking. At predetermined time points, 500 μL of the dialysate was collected. The released DOX was quantified by HPLC.

### Intracellular Uptake of NP-Cy5.5-CRFP

The intracellular uptake of NP-Cy5.5-CRFP was evaluated by flow cytometry and confocal laser scanning microscopy (CLSM). RAW 264.7 cells were incubated with NP-Cy5.5-CRFP for 1, 4, and 7 h. After incubation, the cells were washed with PBS and analyzed by flow cytometry. For CLSM observation, the cells were fixed and stained with a FITC-labeled actin fluorescent probe and DAPI according to the manufacturers’ instructions. The cells were then imaged by CLSM.

### Apoptosis Rate of RAW 264.7 Cells

RAW 264.7 cells were seeded in 6-well plates and treated with LPS, H-151, catechin, or NP-H-CRFP for 24 h. The concentrations of H-151, catechin, and NP-H-CRFP were set at 25 μM, with NP-H-CRFP calculated based on the equivalent H-151 concentration. After treatment, both floating and adherent cells were collected, washed with PBS, and stained with Annexin V-FITC and PI according to the manufacturer’s instructions. The apoptotic cells were analyzed by flow cytometry, and the data were quantified using FlowJo software.

### Cellular ROS Detection

Intracellular ROS levels were detected using DCFH-DA. Briefly, RAW 264.7 cells were seeded in 24-well plates and treated with LPS, H-151, catechin, or NP-H-CRFP for 24 h. H-151, catechin, and NP-H-CRFP were used at 25 μM, with NP-H-CRFP calculated based on the equivalent H-151 concentration. After treatment, the cells were incubated with DCFH-DA in serum-free medium at 37 °C for 30 min in the dark. After washing with PBS, the cells were counterstained with Hoechst and observed by CLSM. In parallel, intracellular ROS fluorescence intensity was analyzed by flow cytometry and quantified using FlowJo software.

### Preparation of Lyophilized Silk Fibroin Powder

Lyophilized silk fibroin powder was prepared according to a previously reported protocol.

### Preparation of Nanoparticle-Loaded Silk Fibroin Hydrogel

Silk fibroin hydrogels were prepared according to a previously reported procedure with slight modifications. Briefly, lyophilized silk fibroin powder was dissolved and mixed with an HRP/H₂O₂-mediated crosslinking system. NP@H-CRFP was then added and thoroughly mixed to obtain a nanoparticle-containing silk fibroin precursor solution. The precursor solution was allowed to undergo gelation at 60 °C to form NP@H-CRFP-loaded silk fibroin hydrogel (Gel-NP). Blank hydrogel (Gel), H-151-loaded hydrogel (Gel-H151), and CRFP-loaded hydrogel (Gel-CRFP) were prepared using the same procedure.

### Preparation of Composite Hydrogel Meshes

Composite hydrogel meshes, including SFM@Gel, SFM@Gel-H-151, SFM@Gel-CRFP, and SFM@Gel-NP, were fabricated using a sandwich-molding strategy. Briefly, a silk fibroin mesh mesh (SFM) was placed in the center of a 20 × 10 × 5 mm³ mold, serving as the intermediate supporting layer and dividing the mold into two symmetric upper and lower chambers. The corresponding silk fibroin hydrogel precursor solutions were then injected into both chambers using a syringe, allowing the precursor solution to fully cover and infiltrate the SFM surface. After gelation at 60 °C, composite hydrogel meshes with a “gel–mesh–gel” sandwich architecture were obtained. Specifically, blank Gel, Gel-H151, Gel-CRFP, or Gel-NP was loaded onto both sides of the SFM to obtain SFM@Gel, SFM@Gel-H151, SFM@Gel-CRFP, and SFM@Gel-NP, respectively.

### Scanning Electron Microscopy

The surface morphology and cross-sectional structure of SFM and SFM@Gel-NP were examined by scanning electron microscopy (SEM). Samples were first frozen at −80 °C and lyophilized for 24 h. For cross-sectional observation, the freeze-dried samples were fractured in liquid nitrogen to expose the internal structure. The samples were then mounted on conductive adhesive tape and sputter-coated with gold. SEM images were acquired at an accelerating voltage. The morphology of SFM and SFM@Gel-NP was compared to evaluate hydrogel loading on the SFM and the interfacial integration between the Gel-NP layer and SFM.

### Rheological Characterization

The apparent rheological properties of SFM@Gel-NP were measured using an ARES-G2 rotational rheometer equipped with a 25.0 mm parallel-plate geometry (TA Instruments, New Castle, DE, USA). Before measurement, the preformed sample was placed at the center of the lower plate, the gap was adjusted according to the sample thickness, and excess material at the edge was removed. All tests were performed at 25 ± 0.1 °C, and silicone oil was applied around the sample edge to minimize water evaporation. Strain-sweep measurements were first conducted at a fixed angular frequency of 6.28 rad s⁻¹ over a strain range of 0.001–10% to determine the linear viscoelastic region. A strain amplitude of 0.1% was then selected for frequency-sweep and time-sweep measurements. The frequency sweep was performed from 0.1 to 100 rad s⁻¹, and the time sweep was conducted at 6.28 rad s⁻¹ for 200 s.

### Mechanical Characterization

Mechanical properties were evaluated using an IPBF-300 in situ mechanical testing system (Tianjin Kaier Measurement and Control Testing System Co., Ltd.). Standardized specimens (1 cm × 2 cm) were clamped with a gauge length of 50 mm and subjected to uniaxial tensile loading at a constant displacement rate of 0.05 mm/s until failure or a maximum displacement of 20 mm was reached. Stress–strain curves were generated from the synchronously recorded force–displacement data.

The ultimate tensile strength (UTS) was defined as the maximum force sustained per unit width (N/cm) prior to fracture under uniaxial tension. n = 3 independent samples were analyzed for each group.

### Swelling Behavior

The swelling behavior of different meshes was evaluated by a gravimetric method. Freeze-dried SFM, SFM@Gel, SFM@Gel-H151, SFM@Gel-CRFP, and SFM@Gel-NP samples were weighed to obtain the initial dry weight W₀. The samples were then immersed in PBS. At predetermined time points, the samples were removed, and excess surface water was gently blotted with filter paper. The swollen weight was recorded as Wt. The swelling ratio was calculated using the following equation:

Swelling ratio (%) = (Wt − W₀) / W₀ × 100%

n = 3 independent samples were analyzed for each group.

### In Vitro Degradation

The enzymatic degradation of different meshes was evaluated by a gravimetric method. Briefly, freeze-dried samples were weighed as the initial dry weight W₀ and then immersed in PBS containing 10 mg mL⁻¹ α-trypsin at 37 °C. At predetermined time points, the samples were collected, gently rinsed with deionized water, lyophilized, and weighed as the remaining dry weight Wt. The degradation ratio was calculated as follows:

Degradation ratio (%) = (W₀ − Wt) / W₀ × 100%

n = 3 independent samples were analyzed for each group.

### Cell viability assessment

Rat-1 cells were seeded in 96-well plates (Corning, USA) at a density of 2 x 10^3^ cells/well and cultured at 37℃with 5% CO_2_. Cell viability was assessed using the CCK-8 assay. After treatment with the drug for 1, 3 and 5 days, 10µL of CCK-8 reagent was added per well (n=6 biological replicates per group), incubated for 2 h, and absorbance was measured at 450 nm.

### Ethical statement and preclinical model

All animal experiments were approved by the Institutional Animal Care and Use Committee (IACUC) of Peking University People’s Hospital (Approval No. 2024PHE003). Female Sprague-Dawley rats (7 weeks old, 200 ± 20 g; Charles River Laboratories, China) were acclimatized to specific pathogen-free (SPF) conditions for 7 days prior to surgery. The rats were randomly divided into the following groups: SFM, SFM@Gel, SFM@Gel-H151, SFM@Gel-CRFP, and SFM@Gel-NP, with 5 rats per group. Anesthesia was induced by inhalation of 5% isoflurane/95% O_2_ at a flow rate of 5L/min and maintained at 2.5%. After shaving and iodine disinfection, bilateral partial-thickness defects (1×2 cm^2^) were created by excising the external and internal oblique muscles 0.5 cm lateral to the linea alba. Meshes were implanted using an intra-abdominal overlay technique, secured tension-free to the surrounding fascia with interrupted 5-0 non-absorbable sutures. In the control group, the defect was closed directly with sutures without mesh implantation. Postoperatively, animals were housed individually and observed routinely.

### Reactive oxygen species (ROS)

ROS levels in tissues were detected using the DCFH-DA fluorescent probe. Mesh-tissue specimens were collected at designated endpoints, embedded in OCT compound, and immediately sectioned at 8 μm thickness using a cryostat. Sections were washed three times with PBS and incubated with 10 μM DCFH-DA working solution for 30 min at 37°C in the dark. After incubation, sections were washed three times with PBS to remove unbound probe. Nuclei were counterstained with DAPI for 5 min. After mounting, images were acquired using a confocal microscope (Leica, Germany). DCF fluorescence signals were semi-quantitatively analyzed using ImageJ software, with five random fields quantified per sample.

### Enzyme-linked immunosorbent assay (ELISA)

Blood samples were collected from rats via the tail vein on postoperative days 1-3 and day 7. After standing at room temperature for 30 min to allow clotting, serum was separated by centrifugation at 3000 rpm for 15 min at 4℃ and assayed immediately. The absorbance at 450 nm was measured, and cytokine concentrations were calculated by interpolation from standard curves. Each sample was assayed in triplicate.

### Transcriptome sequencing and analysis

Total RNA was extracted from mesh-tissue complexes (n=3 per group) using TRIzol (Invitrogen, USA). Libraries were prepared from enriched mRNA, quality-checked (Qsep-400), followed by paired-end 150bp sequencing on the BGI DNBSEQ-T7 platform. Differential gene expression was analyzed using DESeq2 (v1.26.0), with |log_2_(fold change)| > 2 and adjusted *P* < 0.01 defining significant differentially expressed genes (DEGs). Functional enrichment was assessed *via* GO, KEGG and GSEA.

### Western blot analysis

Protein extraction was performed using RIPA reagent (Solarbio, China) containing 1x phosphatase inhibitor cocktail (Solarbio, China). The lysate was collected, and a BCA assay with BSA as standard was adopted to determine protein concentration. Equal amounts of protein were separated by SDS-PAGE and transferred to PVDF membranes at 250V. After blocking with 5% non-fat milk in TBST for 2 h at room temperature, membranes were incubated overnight at 4℃ with primary antibodies, followed by incubation with secondary antibodies for 1 h at room temperature. Signals were detected by chemiluminescence and quantified using ImageJ.

### Immunohistochemistry (IHC) staining

At designated endpoints, rats were euthanized and the mesh-tissue complexes were excised. Mesh-tissue samples (n=5) were fixed, dehydrated, cleared, embedded in paraffin wax, and then cut in 4 μm-thick serial sections. After morphological evaluation by hematoxylin-eosin staining to assess tissue integrity and inflammatory infiltration, IHC was performed. Endogenous peroxidase activity was blocked with 3% hydrogen peroxide solution for 15 min at room temperature. To exclude non-specific binding, sections were incubated with 10% normal goat serum for 30 min at room temperature. Subsequently, sections were incubated overnight at 4℃ with primary antibodies against IL-1β and IL-6, followed by incubation with a horseradish peroxidase (HRP)-conjugated secondary antibody for 1 h at room temperature.

Antibody binding sites were visualized using the DAB chromogen (Servicebio, China). Sections were counterstained with hematoxylin, dehydrated, cleared, and mounted with neutral balsam. Five random fields per section were quantified using ImageJ software, and the percentage of positive area was calculated.

### Histopathological Analysis

Tissue samples of mesh-tissue complexes from rats (n=6) were fixed in 4% neutral buffered formalin for 48 h, embedded in paraffin, and sectioned at 4 μm thickness. Sections were stained with hematoxylin and eosin (H&E) to evaluate inflammatory cell infiltration. Inflammatory reactions were semi-quantitatively assessed using the validated Matthews-Valentin scoring system. Masson’s trichrome staining was performed to visualize collagen deposition, with collagen fibers appearing blue. Five random high-power fields per section were quantified using ImageJ software to calculate the collagen area percentage.

For collagen typing, sections were stained with 0.1% Sirius Red solution for 1 h at room temperature and observed under polarized light. Type I collagen appeared as bright red or orange-yellow, strongly birefringent fibers, while type III collagen appeared as green, weakly birefringent fibers. Five random fields per section were captured, and the areas of type I and type III collagen were quantified using ImageJ software. The type I/III collagen ratio was subsequently calculated.

### Immunofluorescence staining

Paraffin sections were deparaffinized, rehydrated, and subjected to antigen retrieval by microwave heating in sodium citrate buffer. Sections were blocked with 10% normal goat serum containing 0.3% Triton X-100 for 1 h at room temperature. After blocking, sections were incubated overnight at 4°C in a humidified chamber with the following primary antibodies against CD68, CD86 (an M1 macrophage marker), and CD206 (an M2 macrophage marker). After three washes with PBS, sections were incubated with corresponding fluorescent secondary antibodies for 1 h at room temperature in the dark. Following PBS washes, sections were mounted with anti-fade mounting medium containing DAPI. Images were acquired using a confocal microscope (Leica, Germany). Five random high-power fields per section were captured, and the numbers of CD68^+^ cells (total macrophages), CD86^+^ cells (M1 type), and CD206^+^ cells (M2 type) were counted using ImageJ software. The M1/M2 ratio was subsequently calculated.

### Statistical Analysis

All quantitative data are presented as mean ± standard deviation (SD). Statistical analyses were performed using GraphPad Prism software (version 10.0). For comparisons between two groups, an unpaired Student’s t-test was applied.For comparisons involving three or more groups, one-way analysis of variance (ANOVA) was used, followed by Tukey’s post hoc test for multiple comparisons. Statistical significance was defined as P < 0.05, with the following notation: *P < 0.05, **P < 0.01, ***P < 0.001, ****P < 0.0001.

## Acknowledgements

This work was supported by The National Key R&D Program of China (Grant No. 2023YFC2411203) and The Beijing High-Level Innovation and Entrepreneurship Talent Support Program (Grant No. 20250012).

