## Supplemental Figures for "A Sandwich-Structured Silk Fibroin Mesh with ROS-Responsive and Immunoregulatory Functions for Pelvic Floor Repair"

**Supporting Information**

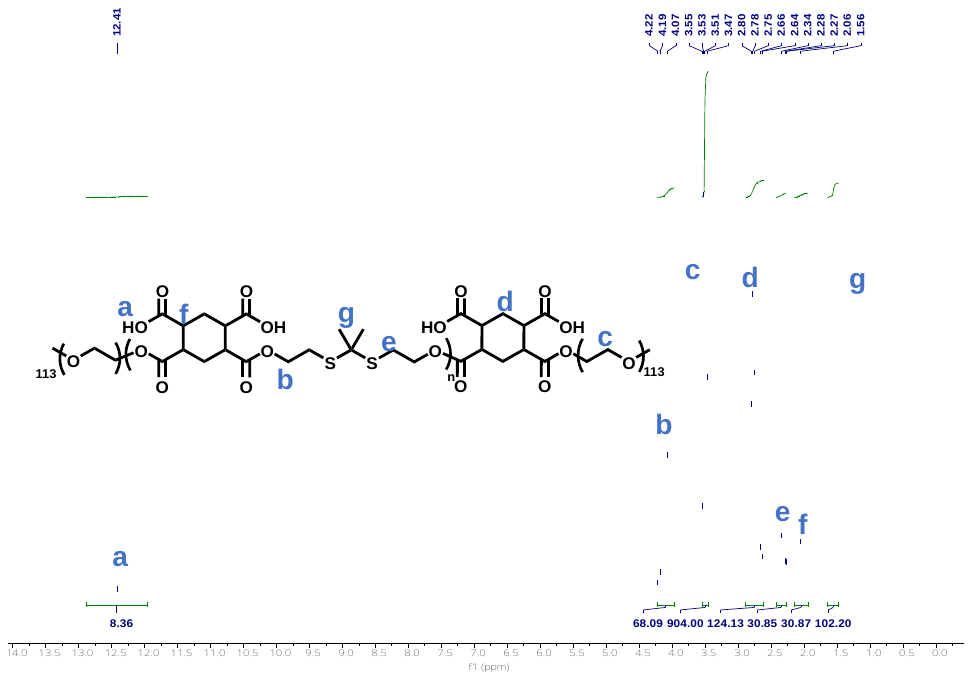

Figure S1 The ^1^H-NMR spectrum of PM077

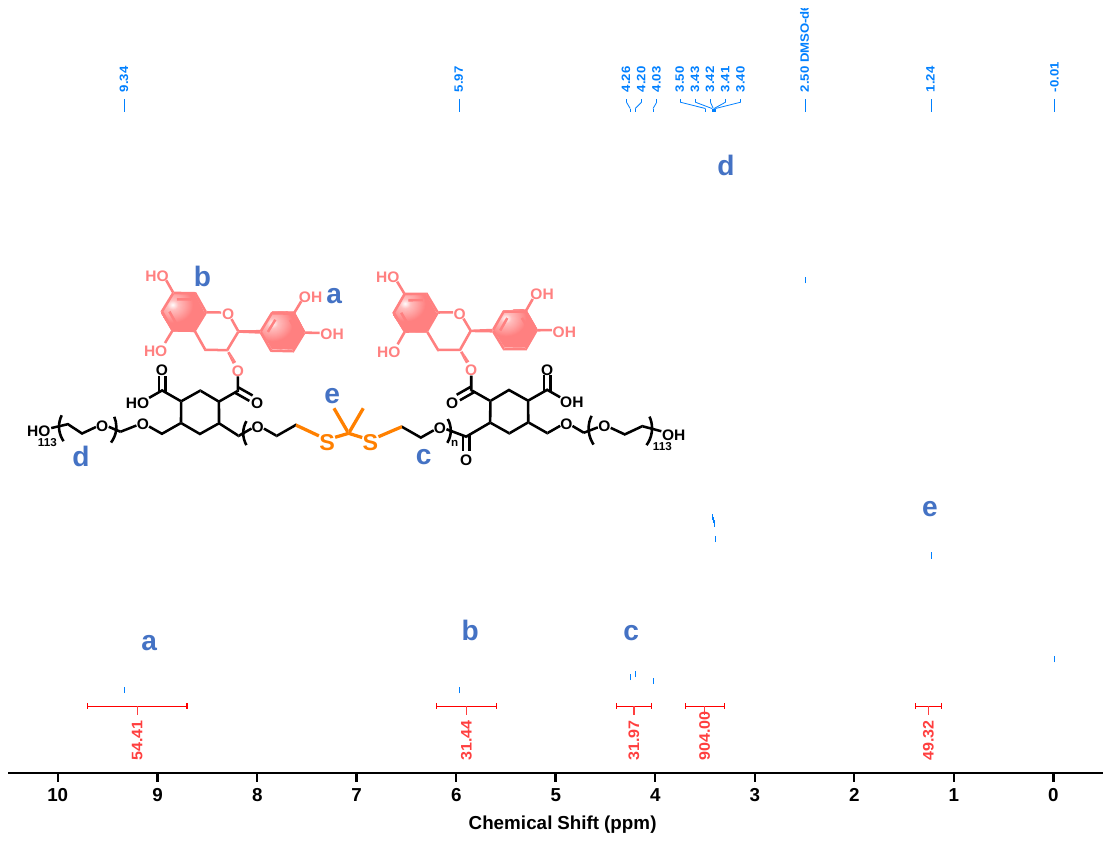

Figure S2 The ^1^H-NMR spectrum of CRFP

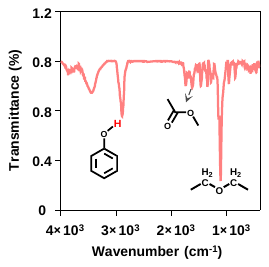

Figure S3 Fourier-transform infrared (FTIR) spectrum of CRFP.

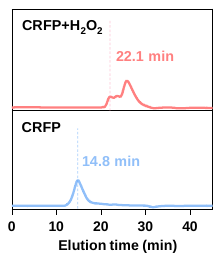

Figure S4 Degradation of CRFP incubation with 10 mM H₂O₂ for 24 h monitored by GPC.

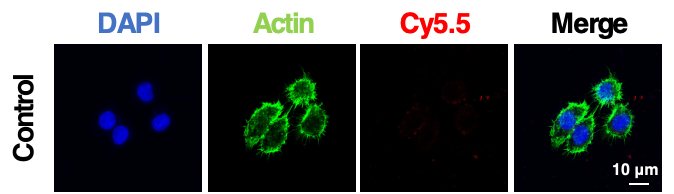

Figure S5 Confocal laser scanning microscopy (CLSM) images of RAW 264.7 cells after incubation with NP-Cy5.5-CRFP for 0 h. Nuclei, cytoskeleton, and NP-Cy5.5-CRFP are shown in blue, green, and red, respectively. Scale bar: 10 μm.

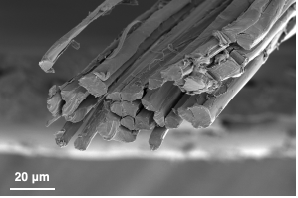

Figure S6 Cross-sectional scanning electron microscopy(SEM) image of SFM. Scale bar: 20 μm.

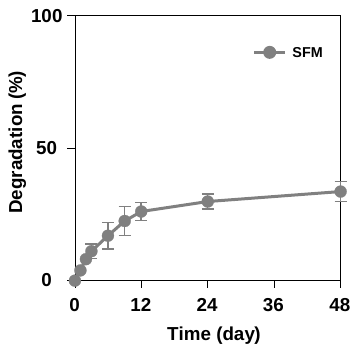

Figure S7 (I) In vitro degradation profiles of SFM.

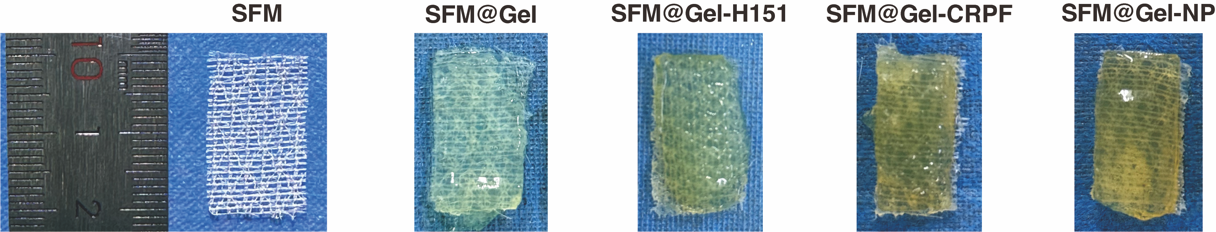

Figure S8. The schematic illustrates the five experimental mesh groups, arranged from left to right as follows: blank SFM (control), SFM coated with blank hydrogel (SFM@Gel), SFM coated with H-151-loaded hydrogel (SFM@Gel-H151), SFM coated with nanoparticle-loaded hydrogel (SFM@Gel-CRFP), and SFM coated with dual-drug-loaded hydrogel (SFM@Gel-NP).

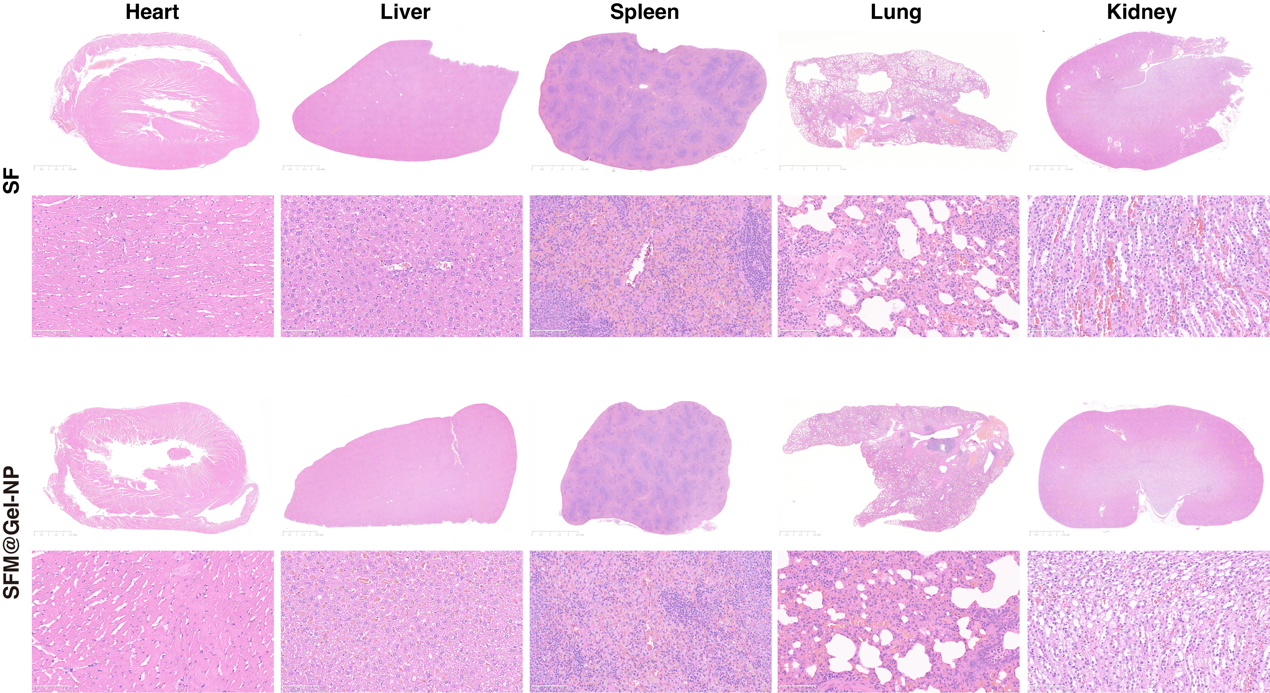

Figure S9. Histopathological assessment of major organs confirms systemic biosafety.

Representative H&E-stained images of the heart, liver, spleen, lung, and kidney tissues harvested from rats in the SFMgroup and the SFM@Gel-NP group at 1 week post-implantation. Scale bars, 2.5 mm.

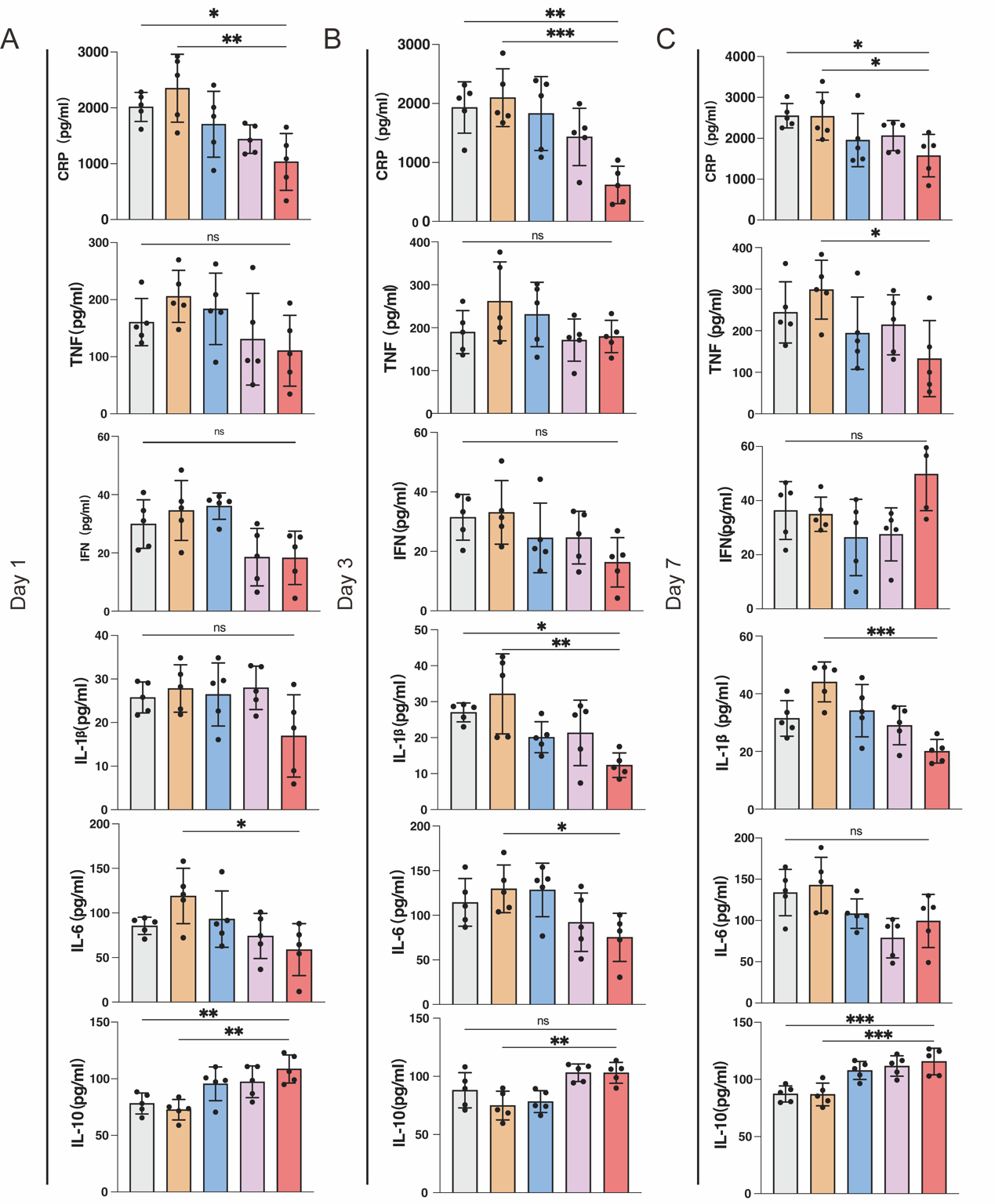

Figure S10. Temporal profile of serum cytokine levels post-implantation.

Serum concentrations of pro-inflammatory cytokines (IL-1β, CRP, IL-6, IFN-γ, TNF-α) and the anti-inflammatory cytokine IL-10 were measured at postoperative days 1, 3, and 7. Data are presented as mean ± SD (n = 5). *P < 0.05, **P < 0.01, ***P < 0.005, ****P < 0.001.

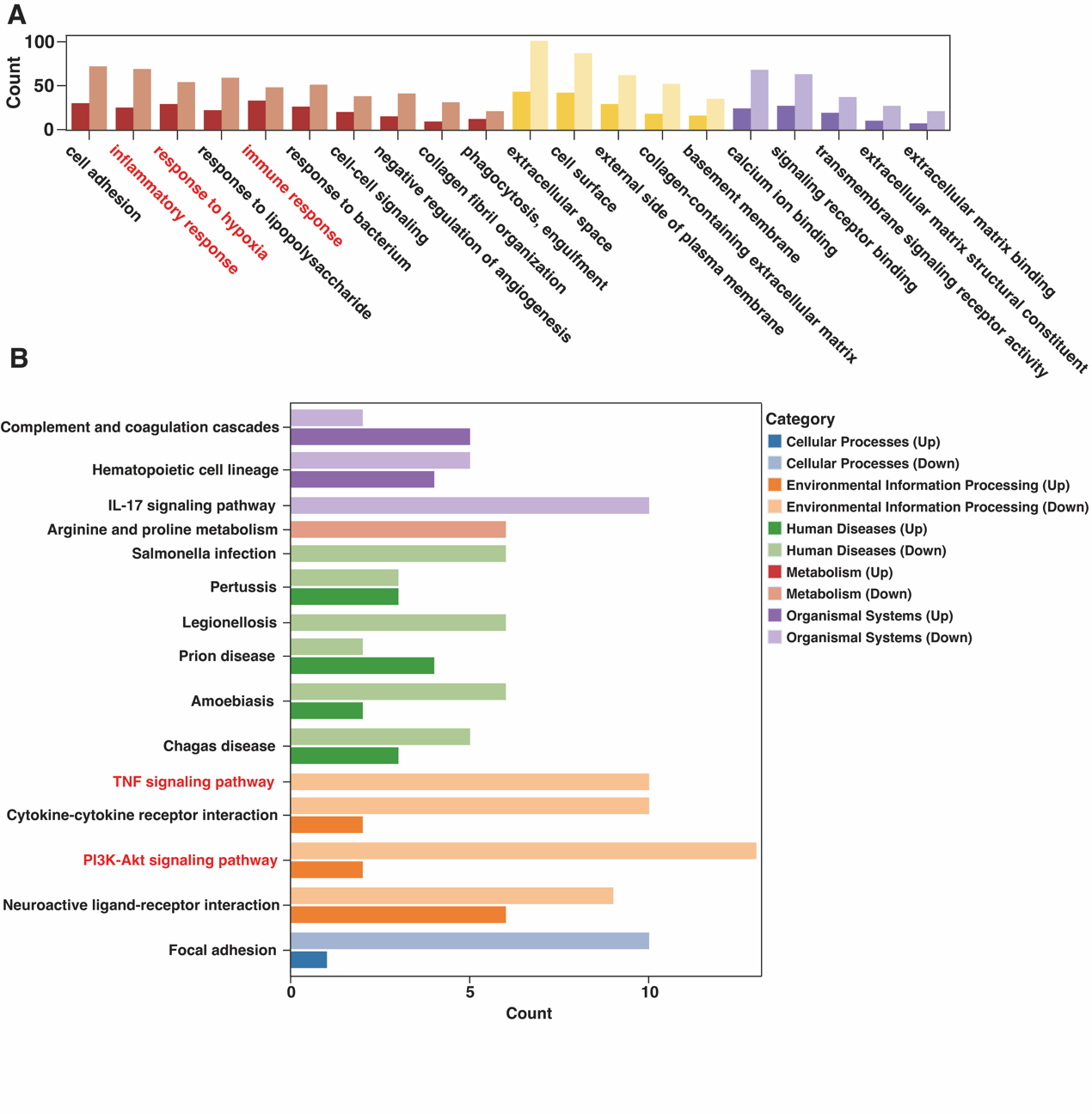

Figure S11. Bioinformatics enrichment analysis of transcriptomic data.

(A) Gene Ontology (GO) enrichment analysis of biological processes for the downregulated differentially expressed genes (DEGs) in the SFM@Gel-NP group compared to the SFM group.

(B) Kyoto Encyclopedia of Genes and Genomes (KEGG) pathway enrichment analysis of the downregulated DEGs. The PI3K-Akt and TNF signaling pathways, key upstream regulators of NF-κB, are highlighted.

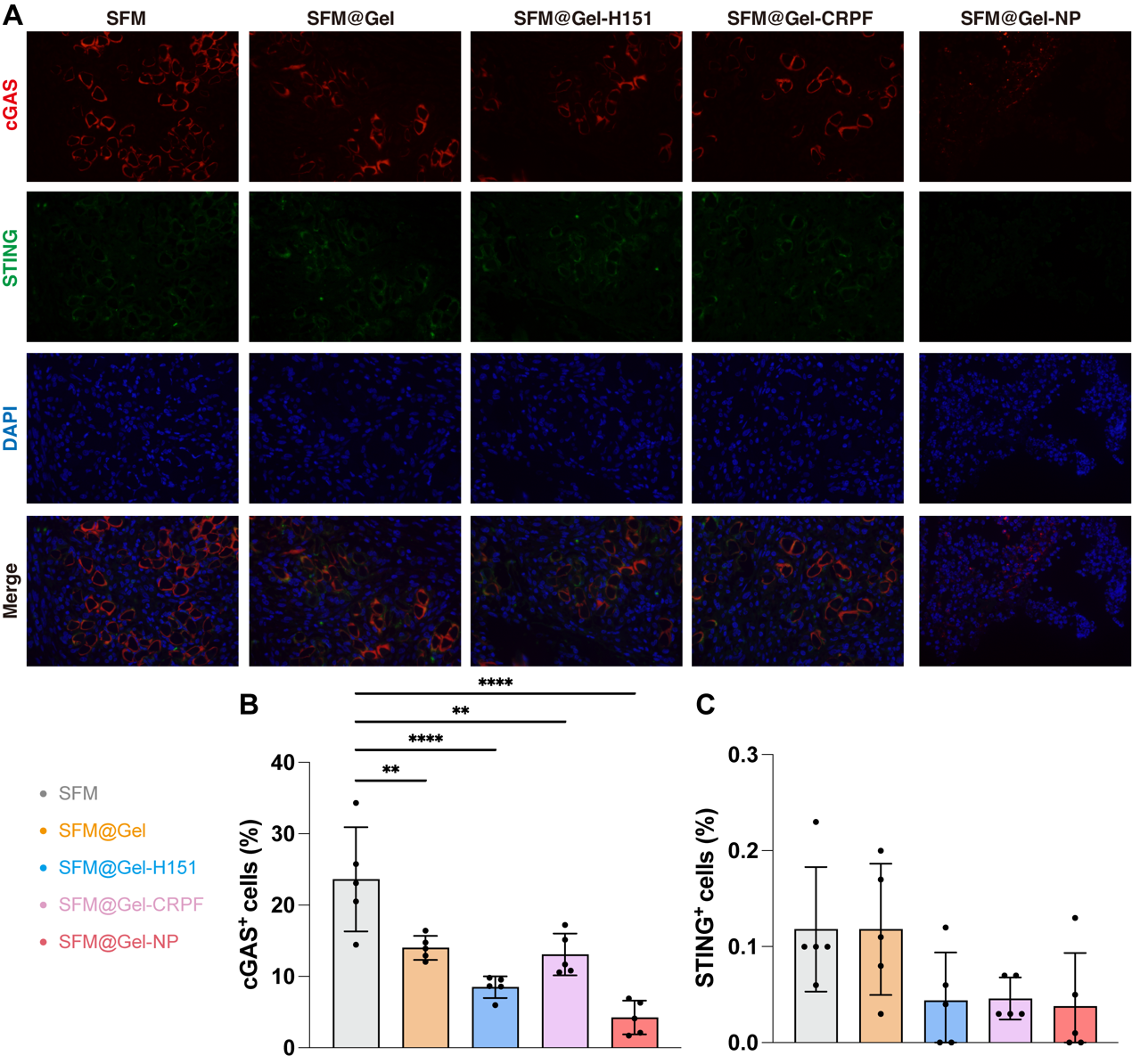

Figure S12. Immunofluorescence analysis of cGAS and STING expression in peri-implant tissues.

(A) Representative immunofluorescence images showing expression of cGAS (red) and STING (green) in tissues surrounding the implant at 1-week post-surgery. Nuclei are counterstained with DAPI (blue). Scale bar, 50 µm.

(B, C) Quantitative analysis of the fluorescence intensity for cGAS (B) and STING (C). Data are presented as mean ± SD (n = 5). *P < 0.05, **P < 0.01, ***P < 0.005, ****P < 0.001.
